# Studying functional brain networks from dry electrode EEG set during music and resting states in neurodevelopment disorder

**DOI:** 10.1101/759738

**Authors:** Ekansh Sareen, Anubha Gupta, Rohit Verma, G. Krishnaveni Achary, Blessin Varkey

## Abstract

There has been an emerging interest in the study of functional brain networks in cognitive neuroscience in order to better understand brain responses to different stimuli. Such studies can help in understanding brain connectivity alterations that arise in neurodevelopmental disorders such as intellectual disability (ID). This research contributes to this body of knowledge by studying alterations in brain connectivity in ID compared to the typically developing controls (TDC). Electroencephalography (EEG) data of subjects with ID and TDC is collected through limited channel dry electrode system. Data was analyzed for the auditory and rest state processing along with the study of intra-network connectivity of the brain via clustering coefficients. Research findings indicate evidences for links between the sensory deficits and social impairment in ID individuals.

## I. INTRODUCTION

NEURODEVELOPMENT refers to the development of neurological pathways that influence the performance and functioning of brain in activities such as social skills, memory, attention, etc. Significant and continued disruption to these dynamically inter-related processes through environmental and genetic risks may lead to neurodevelopmental disorders (NDD) and disability [1]. NDD are prominently linked with the impairment in growth and development of central nervous system during early developmental stages [2]. They are associated with a wider range of functionally-diagnosed conditions such as intellectual disability (ID) [3], autism, cerebral palsy, or attention-deficit/hyperactivity disorder (ADHD) as well as medically diagnosed conditions like Down syndrome, Fragile X syndrome or fetal alcohol syndrome [4]. Thus, there is a need for more focused research to provide a wide array of clinical and social supportive services around the needs of such subjects and their families [5].

The focus of this study is on intellectually disabled (ID) population. ID subjects are generally identified by adaptive, cognitive, and social skill deficits [6], and are often accompanied by challenging behavior such as an increased risk of injury [7]. Despite considerable research in exploring psychiatric and social morbidity of ID, little is understood about its association with brain development and organization. This has prompted researchers to adopt different neuroimaging and brain network extraction methods to understand the structural and functional connectivity in response to specific activities.

Structural Connectivity (SC) explores the anatomical connections between brain regions, whereas Functional Connectivity (FC) looks at the interaction between brain regions that aligns with either cognitive performance or brain’s default state [8]. FC can be measured using statistical methods such as correlation, phase coherence, and covariance [9]. This helps in understanding the synchronization of inter-related activities among brain regions via the construction of functional brain networks (fBNs). These fBNs play an important role in understanding the cognitive behaviour of brain. For example, FC has been helpful in extracting motor [10], visual [11], language [12], default mode [13] and attention [14] networks.

In the field of neuroimaging, brain connectivity is predominantly studied in response to certain stimuli, i.e., while the subject is engaged in a task. A lot of research has been done in exploring brain connectivity by examining stimulus-generated brain responses when engaged in cognitive tasks. Music perception is one such task that is known to generate cognitive and perceptive responses linked to memory and emotions [15]. Recent studies have suggested significantly enhanced phase synchrony while listening to music as compared to resting condition in healthy subjects [16], [17]. Related studies have also observed that there exist different degrees of functional cooperation between two adjacent brain regions in several musical perception tasks [17]. From a graph theoretical point of view, musical perception depicts increased FC and enhanced small-world network organization.

To explore neurobiology of brain disorders such as ID, studies have adopted different neuroimaging techniques, say diffusion tensor imaging (DTI), functional magnetic resonance imaging (fMRI), electroencephalography (EEG) and magnetoencephalography (MEG) [9]. Use of different data modalities can provide complimentary views on brain connectivity and its variation. Although EEG has better temporal resolution and is more accessible with reference to cost and ease compared to fMRI, it is not utilized actively to extract fBNs, while a plethora of research studies have made use of fMRI for the same. EEG also holds a practical and theoretical advantage in elucidating clinical biomarkers [18]. In fact, it is one of the most cost-effective and clinically-available modalities for mass recording and monitoring treatment outcomes [18], [19]. Recent studies are using EEG to study cognitive load in NDD like Autism [20]. This study is also aimed at exploring EEG data modality for providing an understanding of brain connectivity in ID versus typically developing control (TDC) individuals.

Since graph-theoretic and network science measures are used to assess communication efficiency and organization of networks, they are extensively used to study fBN organization [21]. This study is aimed at finding communities with densely connected set of nodes, but with inter-community connections being scanty in the network. Research has suggested that these community structures indicate strong inter-neuron connections that contribute to functional responses within the brain. Studies have suggested that the prominent network science measures like shorter path length and higher clustering are linked with better cognitive abilities, while deviant brain topologies may be indictors of neuropsychiatric disorders [22]. Thus, exploring fBN organization through the lens of graph theoretic models in neuropsychiatric disorder such as ID is of great relevance.

Advancements in brain network analysis has entailed the need for integration of statistical tools with brain network analysis [24]. Generally, a univariate metric is used to summarize and compare networks with simple tests (e.g. t-test, F-test etc.). These approaches are often used because of their simplicity, but they do not account for local connectivity trend within the network [25]. The network-based statistics (NBS) [26] is one of the few methods that accounts for comparison of nodal properties or local connectivity patterns of a network. However, it fails to account for the topological differences in the entire network of a subject or a group. Thus, there is a need of a statistical measure that considers the local connectivity pattern as well as group differences at the network level. In this work, we propose a group level comparison measure which is inspired from the work of *Simpson et al.* [25], where they used permutation-testing framework to compare a group of networks while incorporating topological features inherent in individual networks. On the similar lines, this study proposes a method that incorporates this permutation-testing framework with a graph-theoretic measure called global clustering coefficient for assessment of group differences at the network level.

For ID individuals, identification of brain connectivity patterns in response to stimuli can enable interventions to these subjects that can assist their caregivers, researchers and the society, in filling the educational gaps and providing them with a better lifestyle. This research study provides a framework to explore functional connectivity and brain network organization in ID using EEG. This work tested and verified the working of EMOTIV’s EPOC+ 14-channel EEG machine that is portable and cost-effective device. Also, this device overcomes experimental limitations associated with the clinical population of interest, i.e., people with ID who are not comfortable with the pasting of wet electrode EEG system. This work also explored the brain network organization associated with music apprehension task in two groups, i.e., ID and TDC, and provides cognitive reasoning for the same. Finally, this work provides a statistical framework to differentiate between ID and TDC groups by using the proposed permutation-testing framework.

## II. MATERIALS

### A. Participants

A total of ten typically developing controls (TDC) and seven intellectually challenged participants took part in the study. Both ID and TDC participants were recruited from NaiDisha Centre, Tamana NGO, New Delhi, India. The ID participants were diagnosed under code range of F70-F79 of ICD-10 medical classification list of mental retardation by World Health Organization (WHO). The age of ID participants (all male), ranged from 26 to 31 years (age = 28.66±1.966) with Intelligence Quotient (IQ) from 52 to 68 (mean IQ: 59.167±5.8452) and Social Quotient (SQ) from 57 to 62 (mean SQ: 59.667±2.0656). The age of TDC participants (all male) ranged from 18 to 56 years (age = 33.85±14.89). Approval for the experiment was granted by the Institute’s Ethics Committee and the experiment was performed in accordance with the ethical standards laid down in the 1964 Declaration of Helsinki under the expert guidance of caregivers who work with NDD population. Malin’s Intelligence Scale for Indian Children (MISIC) that is an Indian adaptation of Wechsler’s Intelligence Scale for Children (WISC) and Vineland Social Maturity Scale were the standardized tools used for the evaluation of IQ and SQ of ID population, respectively.

### B. EEG recording and pre-processing

Raw Electroencephalogram (EEG) data was acquired using a 14-channel wireless EEG headset called EPOC by Emotiv [27] with a sampling rate of 128Hz using a subscription-based software Emotiv Xavier. It uses saline-based wet sensors instead of sticky gels which was a distinguishing feature with other similar products available in the market. Fig. 1 depicts the 14 electrodes of Emotiv EPOC relative to 10-20 electrode system. Fig. S1 (Supplementary) depicts the parcellation of brain into Brodmann Areas (BA), while Table-S1 lists BA along with the cognitive functionality of these electrodes.

**Fig. 1.**
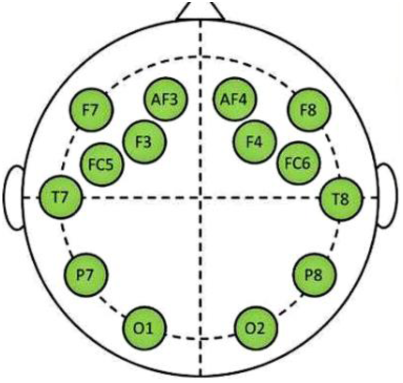
Electrode mapping Source: Adapted from [31]

The subject was seated on a comfortable chair in relaxed position with the wireless headset placed over the head. He was briefed about the experiment having 2 minutes of silence (rest state) and 4-5 minutes of a song (music state). Earphones were plugged in the ears for the auditory stimulus and eyes were closed throughout the session. The subject was also instructed to not engage in stressful thinking during the rest state.

Recorded raw EEG data files were first header-fixed using the EDF Browser software [28] to make these compatible for loading in EEGLAB for filtering, correction, and visualization. Next, the EEG data was filtered and pre-processed using the pipeline implemented using EEGLAB [29] and MATLAB. The pre-processing pipeline included selection of 14 channels of Emotiv and was performed in EEGLAB. This data was filtered using a high pass filter with a lower frequency cut-off set at 0.5Hz in EEGLAB to minimize filtering artifacts at epoch boundaries. A customized channel location file for the EMOTIV device which contains the spatial orientation of its electrodes was created and added. For removing artefacts from the EEG data, Independent Components Analysis (ICA) decomposition of the data using infomax algorithm was performed in EEGLAB. Next, ADJUST plugin [30] was used that automatically computes a set of spatial and temporal features for each of the independent components (ICs). Artefacts are associated with temporal and spatial features as follows: 1) *Eye Blinks:* Spatial Average Difference (SAD) and Temporal Kurtosis (TK), 2) *Vertical eye movements:* SAD and Maximum Epoch Variance (MEV), 3) *Horizontal eye movement:* Spatial Eye Difference (SED) and MEV, and 4) *Generic discontinuities:* Generic Discontinuities Spatial Feature (GDSF) and MEV.

ADJUST displays all ICs along with their features. ICs, that cross the pre-determined thresholds of both spatial and temporal features of any artifact listed above, are highlighted by ADJUST. Finally, the marked artefact components were removed and the new dataset was saved. Here, with 14-channel data, only the worst two components (identified by visual inspection) of all the ADJUST-classified components were rejected. Resulting data was reconstructed to obtain clean, filtered, and artifact-free EEG signal. Fig. 2 provides the flowchart of the steps taken in pre-processing of EEG data. This data was then subdivided according to the rest and music state event markings and was analyzed next.

**Fig. 2.**
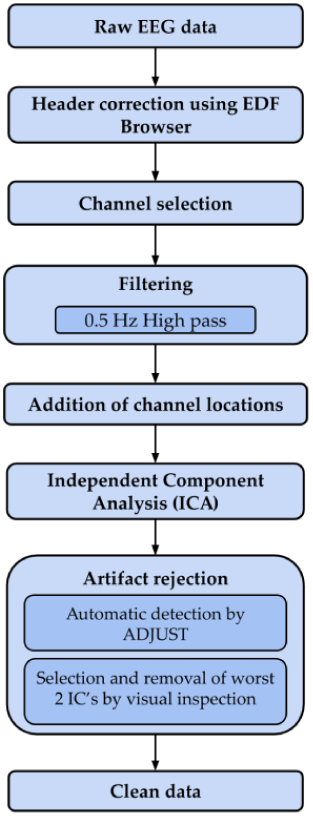
Filtering and pre-processing pipeline

## III. METHODS

### A. Functional connectivity analysis

This study uses partial correlation (PC) to measure statistical coupling between EEG signals. PC coefficients quantify correlation between two sensors, while being controlled for other sensors [32]. Unlike correlation, it prevents false prediction of direct link between two sensors. If *r*_*ij*_ is the Pearson correlation between *‘i’* and *‘j’, r*_*ik*_ is the Pearson correlation between *‘i’* and *‘k’*, and *r*_*jk*_ is the Pearson correlation between *‘j’* and *‘k’*, with *k*=1,2,…*m* as the other sensors to be controlled for, then PC *P*_*ij*_ between sensors *‘i’* and *‘j’* is

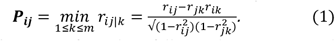

FC analysis was performed as follows: the pre-processed time-series EEG data comprising of *N* = 14 channels was subdivided into *T* number of non-overlapping static windows of *L* samples (say *L*=32) each, resulting in a 2D matrix (or a slice) of size *N×L*. Next, the adjacency matrix was built using the PC of size *N×N*. All time windows of subject ‘*s*’ were stacked to create a 3D tensor **W**_*s*_(*t*) of size *N* × *N* × *T*. Thus, we obtained *S* number of 3D tensors.

### B. Temporal summarization for extracting static connectivity

To study the static FC, connectivity matrix **W**_s_(*t*) of subject *s* was temporally summarized via Higher Order Singular Vector Decomposition (HOSVD), also known as Tucker Decomposition [33], as

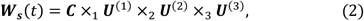

where **C** ∈ ℝ^*N*×*N*×*T*^ is the core tensor and the three orthonormal mode matrices containing the left singular vectors corresponding to each of three modes are depicted by **U**^(1)^ ∈ ℝ^*N*×*N*^, **U**^(2)^ ∈ ℝ^*N*×*N*^, and **U**^(3)^ ∈ ℝ^*T×T*^ or the horizontal, frontal and vertical unitary matrices of **W**_s_(*t*). Unfolding the tensor along the desired mode using singular value decomposition (SVD) results in the corresponding mode matrices. Since the column space of **U**^(3)^ spans the temporal direction, the singular vector corresponding to the highest singular value is utilized for temporal summarization. To achieve this **W**_s_(*t*). was flattened 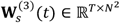 and SVD was applied to find the singular vector of **u**_(3,1)_ that is associated with the largest singular value. This vector captures the most significant contribution in time.

Thus, **u**_(3,1)_ is utilized to carry out weighted temporal summarization of connectivity matrix as

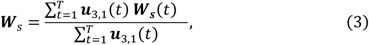

where **W**_s_ ∈ ℝ^*N × N*^ is the static connectivity matrix of subject *‘s’*. Similarly, the summarization was done for all subjects to obtain *S* number of *N×N* connectivity matrices.

### C. Subject summarization for extracting group-level functional connectivity

To study group-level FC, connectivity matrices of all *S* subjects of a particular group (say ID) were stacked together to create a 3D tensor **W** of size *N* × *N* × *S* Similar to the process explained above, subject summarization was carried out after HoSVD [34]. Compared to simple averaging of the connectivity matrices of all subjects, HoSVD based subject summarization accounts for inter-subject variability and carries out weighted summarization. This led to the formation of static group-level adjacency matrix of size *N* × *N*.

### D. Binary graph generation

The subject summarized-weighted 2D adjacency matrix is binarized by applying a threshold to select only the significant connections. Top 30% connections were considered significant and an absolute threshold value was determined for the same. PC values in the connectivity matrix exceeding the threshold were set to *‘*1*’* and rest were set to *‘*0*’*. This resulted in the formation of a binary connection matrix, where a connection or an edge between two nodes exist, if the corresponding entry in the binary connection matrix is *‘*1*’*; otherwise, there is no edge between them.

### E. Graph theoretical metrics

Next, graph metrics were computed from the binary connection matrix of size *N×N*. The network was explored visually. Functional brain networks as well as network metrics were extracted in Gephi [35]. Louvain modularity algorithm with randomized experiments was used to extract unweighted and undirected communities or networks [36]. The quality of partition *P* was quantified using the modularity index *Q* as

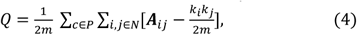

where *‘m’* is the total number of edges, **A** is the adjacency matrix of the network, *k*_*i*_ is the degree of the node *‘i’*. The index *c* runs over the modules of partition *P*, while the indices *‘i’* and *‘j’* span across the *N* nodes of the matrix. Louvain algorithm assigns a modularity class to each node. Networks possessing high modularity can be interpreted as having high inter-community connections and sparse intra-community connections. It is considered to be the best measure of determining quality of partition *P* of a network.

We also explored graph theory metric, namely, clustering coefficient that is considered to be a useful measure to quantify function-structure associations in the brain and helps in the understanding of small-worldness of a network, its community structure and its local efficiency [37]. It is a popular measure for studying functional segregation across different communities and functional aggregation within a community of brain. For example, it measures how likely it is that the two connected nodes are part of a larger highly connected group or degree to which nodes in a network tend to cluster. To quantify this tendency, clustering coefficient was introduced [38]. There are two types of clustering coefficients, local clustering coefficient (LCC) and global clustering coefficient (GCC). LCC is computed for all the nodes in the network and provides the fraction of nodes that are connected to each other. LCC considers local connectivity pattern, while GCC is computed for the entire network and provides average neighborhood connectivity for the entire network. GCC can be computed by two approaches, first by computing the average of LCC over all the nodes. The second approach for computing GCC is by measuring the transitivity of the network, which is the ratio of number of triangles and number of open triads in a network as

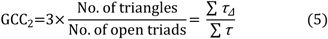

where Σ *τ* gives the total number of open triads and Σ *τ*_*Δ*_ is the subset of these open triads which are triangles. The value of this coefficient ranges between *‘*0*’* and *‘*1*’*. Both the approaches to compute GCC, either by GCC_1_ or by GCC_2_, measure the tendency of edges to form triangles. GCC_2_ places larger weight on high degree nodes as compared to GCC_1_, which can be really useful while studying brain networks.

One of the objectives of the study was to understand whether GCC is a suitable measure to detect group-related differences between TDC and ID groups. Additionally, it was desired to explore as to which of the two approaches of computing GCC is more relevant for such studies. To this end, the 3D tensor of filtered EEG data of dimension *N × L × T × S* was converted into a 4D tensor of dimension *N×N×T×S*, containing the *N×N* connectivity matrices (generated using functional connectivity measure), for all the time windows *T* and for all the subjects *S*. This 4D tensor was then subject-summarized and binarized into a 3D tensor of dimension *N×N×T*, for which the GCC values were computed using both the approaches, to study the evolution of modules and clusters over a period of time. GCC_1_ and GCC_2_ were computed using the Brain Connectivity Toolbox (BCT) [39] over the binary undirected adjacency matrices of dimension *N×N* for all time windows.

### F. Statistical analysis

Brain network analyses and neuroimaging studies have taken a transcendental leap over the past couple of decades. However, for statistical measures, researchers usually resort to common statistical tests that rely on network metric and univariate nodal or edge-based comparisons that ignore the inherent topological properties of the network. This study carries out statistical testing by two methods. The first method is the conventional univariate statistical comparison of results by the non-parametric two-sample *t*-test, i.e., the Wilcoxon rank sum test that does not assume known distributions. This test returns a decision for the null hypothesis that data in two samples are from continuous distributions with equal medians, against the alternative that they are not. This statistical test was applied on the GCC values to examine if GCC values were statistically significantly different in the two groups TDC and ID for both rest and music state. Results from this test provided statistical support for the cognitive and theoretic findings of this work.

Inspired from the permutation testing framework of Simpson et al. [25], this work proposed to statistically test the clustering strength of networks of ID and TDC groups via permutation testing. In permutation testing, the distribution of test statistic under the null hypothesis is generated empirically from the data by permuting data labels. Analysis of GCC, which is the degree to which nodes in a graph tend to form a cluster, provided an insight of the regions of activity in the brain and thus, of the functional brain networks.

## IV. RESULTS AND DISCUSSION

### A. ID and TDC Group-level Network analysis

The inter-group differences between TDC and ID population was studied from the perspective of network analysis and cognitive reasonings associated with nodal properties of the network. A total of four networks were generated for group-level analysis for rest and music states of ID and TDC groups. After the binarization of adjacency matrix to ‘0’ and ‘1’ as described earlier in Section III-D, some nodes were not observed to have connection with any other node and hence, were not available in the final networks (missing nodes). Also, diagonals of the adjacency matrix that denote the connection of a node to itself (self-loop) were not considered.

The fBNs generated for group level analysis of the TDC group, for both rest and music state are depicted in Fig. 3 and Fig. 4. Among a wide array of inter-nodal connections, three network communities were found in both rest and music state albeit with subtle differences in their formation in respective states. The predominance of nodes with respect to their nodal connections was, however, majorly changed while still remaining in the similar network formation in both states.

**Fig. 3.**
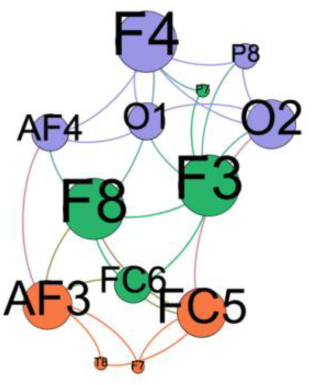
Rest state network for TDC group (Nodes of same color denote one community; three communities are detected in this figure)

**Fig. 4.**
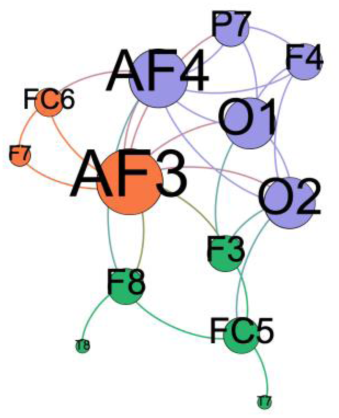
Music state network for TDC group (Nodes of same color denote one community; three communities are detected in this figure)

During the rest state, the first community of nodes had AF3 (BA09), FC5 (BA44), T8 (BA21) and F7 (BA45 and BA47) with a predominant connectivity at AF3 and FC5 nodes. The second community of nodes had F3 (BA08), F8 (BA45 and BA47), FC6 (BA06) and P7 (BA19 and BA37) with predominant connectivity at F3 and F8 nodes. The third community of nodes had F4 (BA08), AF4 (BA09), P8 (BA19 and BA37), O1 (BA18) and O2 (BA18) with predominant connectivity at F4 and O2 nodes. During the music state, three slightly different network communities were observed for TDC group. The first community of nodes had AF3 (BA09), FC6 (BA06) and F7 (BA45 and BA47) with a predominant connectivity at AF3 node. The second community of nodes had F3 (BA08), F8 (BA45 and BA47), FC5 (BA44), T7 (BA21) and T8 (BA21). The third community of nodes had F4 (BA08), AF4 (BA09), P7 (BA19 and BA37), O1 (BA18) and O2 (BA18) with a predominant connectivity at AF4, O1 and O2 nodes.

During the rest-state, bilateral frontal cortex showed dominant activity with minimal to no connection with temporo-parietal regions. The dominant connections were at F3 and F4 nodes that are considered relevant for secondary motor response, motor planning, imaging and learning, auditory imagery, working memory and visuo-motor response. Such strong frontal behaviour of this community can be explained by subject’s engagement in working memory, memory retrieval and mental imaging which happens during the rest state, when the mind is engaged in random thought processes. Negligible activation on temporal sites can be explained by no auditory stimulus and complete silence during the rest state. But during the music state, temporal cortex had activated to significant proportions. Such strong temporal behaviour of this community can be explained by the auditory stimulus provided to the subjects, which resulted in significant activation at temporal sites. During the music state, temporal nodes had increased connections to the respective ipsilateral frontal regions with increased connectivity at the dorsolateral prefrontal cortex (AF3 and AF4), which is a major seat for language comprehension and expression, working memory, memory retrieval and encoding, selective attention to sounds, executive planning, auditory imagery and emotional stimuli. This could reflect the activation of fronto-temporal circuitry during music perception.

The interconnection of nodes was not just lateralized to a hemisphere, but had inter-hemispheric connections too. This implies the importance of interhemispheric neurons forming communications with different cortical areas. The presence of distinct occipito-parietal network in both states further validates the presence of different networks observed in the current study. The activation of occipital network can be due to visual mental imaging that goes on while at rest with closed eyes and can be explained by subject’s engagement in semantic processing and memory recalling due to the given auditory stimulus in music state. It was also observed that networks were more intermingled at rest but became slightly more distinctive but yet incorporated with music.

From the above network analysis of TDC group for both rest and music state, we can conclude that the generated functional networks are consistent in nature, as their network properties stand in complete correlation not only with the cognitive reasoning suggested by related literature but also with the experimental design. Moreover, this conclusion also validates the efficiency of the 14-electrode wireless EEG device in recording electrophysiological brain activity, which can be used to generate consistent functional brain networks.

After validating the recording device and establishing the consistency of generated functional brain network, the functional networks generated for group-level analysis of the ID group for both rest and music state were explored. The functional networks generated for group level analysis of the ID group for both rest and music states are depicted in Fig. 5 and Fig. 6, respectively. Among a wide array of inter-nodal connections, two network communities were found in the rest state, while four communities were found in the music state.

**Fig. 5.**
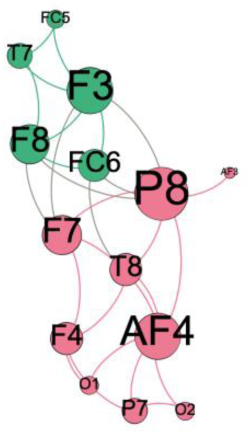
Rest state network for ID group (Nodes of same color denote one community; two communities are detected in this figure)

**Fig. 6.**
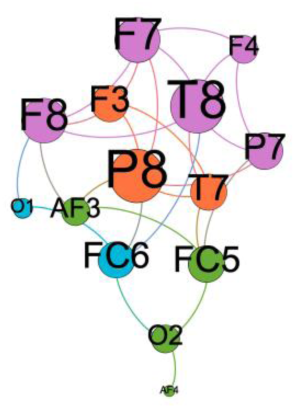
Music state network for ID group (Nodes of same color denote one community; four communities are detected in this figure)

During the rest state, the first community of nodes had AF3 (BA09), AF4 (BA09), FC5 (BA44), F4 (BA08), P7 (BA19 and BA37), P8 (BA19 and BA37), F7 (BA45 and BA47), T8 (BA21), O1 (BA18) and O2 (BA18) with a predominant connectivity at AF4 and P8 nodes. The second community of nodes had F3 (BA08), F8 (BA45 and BA47), FC5 (BA44), FC6 (BA06) and T7 (BA21) with predominant connectivity at F3 node. The internodal connection was observed to be sparse in ID group as compared to the TDC group during rest state suggesting poorer inter- and intra-hemispheric connection in ID group. Overall, during the rest state, there was a dominance of parietal activity of the dominant hemisphere in ID group in comparison to the bilateral frontal dominance as seen in TDC group suggesting poorer attention and lesser involvement in executive planning in ID group participants.

During the music state, four network communities were observed for ID group. The first community of nodes had AF3 (BA09), AF4 (BA09), FC5 (BA44) and O2 (BA18) with predominant connectivity of FC5 node. The second community of nodes had FC6 (BA06) and O1 (BA18) with predominant connectivity of FC6 node. The third community of nodes had P8 (BA19 and BA37), F3 (BA08) and T7 (BA21) with predominant connectivity of P8 node. The fourth network was composed of F4 (BA08), F7 (BA45 and BA47), F8 (BA45 and BA47), P7 (BA19 and BA37) and T8 (BA21) with predominant connectivity of T8, F7 and F8 nodes. Segregation of networks in music state in comparison to rest state networks suggests that parallel circuits are formed when stimuli such as music is provided to ID participants. This may reflect poorer integration of cortical circuitry requiring more neuronal efforts for the processing of audio stimuli. There was a predominance of P8 connections (representing the inferior temporal gyrus of dominant hemisphere) in both music and rest states in ID group participants suggests more importance is provided to environmental structures such as face and body recognition. The enhanced activity at temporal regions (T8 more than T7) during the music state could be understood in the context of their involvement in the perception of auditory stimuli during listening music.

During the music state, the predominance of frontal connections was not observed in ID group as in the TDC group. Rather, an overall predominance was observed at the temporo-parietal region. Also, the ID group lacked in the activation of the predominant fronto-temporal circuits as observed in the TDC group during the music state. Moreover, unlike the distinctive activity at the dorsolateral prefrontal cortex circuitry in TDC group, the participants in the ID group displayed poorer involvement of this region reflecting lesser involvement in executive processing of music. The predominant connections of T8 to F7 and F8 and belonging to a separate network further suggests music enjoyment rather than appropriate audio processing of the stimuli in context of executive functioning. The separate occipito-parietal community network observed in TDC group was not observed in ID group in either state supporting poorer networking circuits in these groups of participants.

With the above network analysis, it was concluded that the generated functional networks were consistent with reference to the cognitive reasonings associated with the structural property of networks.

### B. Statistical analysis of network parameter

In general, it is known that there is less coordination between brain regions in NDD. This study attempted to validate this via statistical analysis of the GCC as a network parameter. Results of Wilcoxon ranksum test suggested the following: 1) for GCC-1, significant differences were observed between TDC and ID classes in the rest state (*p*=3.29e-51, Table S2) and, between TDC and ID classes in the music state (*p*=7.84e-106, Table S2), 2) for GCC-2, again significant differences were observed between TDC and ID classes in the rest state (*p*=3.10E-86, Table S2) and, between TDC and ID classes in the music state (*p*=1.88e-184, Table S2) and, 3) much lower *p*-values were observed while using GCC-2 as a parameter compared to GCC-1, indicating that GCC-2 is a better measure of the two because it is more efficient in capturing differences between the two groups in group-level analysis.

Apart from comparing the two groups in rest and music states, data from rest and music states of both the groups were also analyzed vis-à-vis each other. Results suggested the following: 1) for GCC-1, no significant differences were observed between rest and music classes for the TDC group (*p*=0.0422, Table S3) and, rest and music classes for the ID group (*p*=0.1232, Table S3), 2) for GCC-2, again no significant differences were observed between rest and music classes for TDC group (*p*=0.0237, Table S3) and, rest and music classes for ID group (*p*=0.0231, Table S3). Fig. 7 and 8 depict boxplots for the visual comparison of the two groups for both GCC-1 and GCC-2 values in rest and music states from which it was concluded that the two groups are indeed different.

**Fig. 7.**
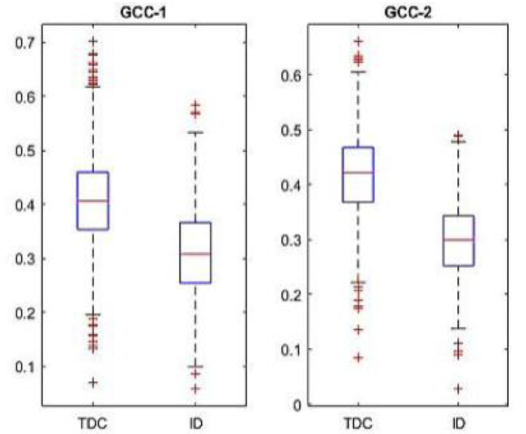
Boxplots for TDC vs ID rest state

**Fig. 8.**
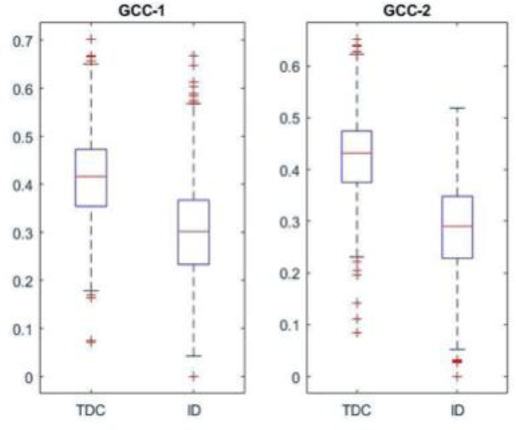
Boxplots for TDC vs ID music state

### C. Permutation testing framework

In the proposed permutation-testing framework, GCC through transitivity measure (GCC-2) was used as the test statistic. This study had seven subjects in the ID group (G1) and ten subjects in the TDC group (G2). For the sake of uniformity in the number of subjects considered, 7 subjects from the TDC group were randomly selected. First, subjects in the dataset were assigned an ‘id’ called as the original label (refer to Fig. 9) and GCC-2 was computed as the test statistic for all the subjects in G1 and G2 groups. Next, ratio of mean of GCC-2 values of G1 to that of G2, denoted by R_gcc_, was computed as depicted in Fig. 9.

**Fig. 9.**
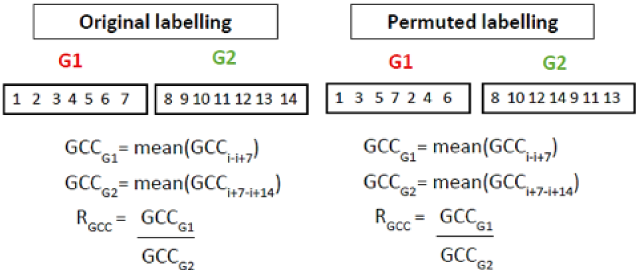
Figure shows labels of two groups G1 and G2, each with 7 subjects. Left: depicts the original labelling of subjects and Right: depicts one example of permuted labelling of subjects

This R_gcc_ value corresponds to the original label. The labels were then permuted (randomly shuffled) and the R_gcc_ values were again computed for all the set of combinations, resulting in *K* number of R_gcc_ values, where *K* is the number of times labels were permuted. After generating *K* number of R_gcc_ values, their histogram was plotted to see the empirical distribution. If the R_gcc_ value computed using the original labels lies beyond 0.05 or lesser probability region of the empirical distribution generated using permutation testing framework, it is said to be significant.

The permutations were done 300,000 times and the empirical distribution of R_gcc_ values was plotted for the music state data of both the groups. Fig. 10 depicts this distribution and also the position of R_gcc_ value calculated using the original labels i.e. observed R_gcc_. From this figure, it was concluded that the functional brain networks of the two groups under study, i.e., the TDC and ID groups, for music state data, were indeed statistically different. Similar procedure was repeated on the rest data. Fig. 11 depicts this distribution and also the position of the R_gcc_ value calculated using the original labels.

**Fig. 10.**
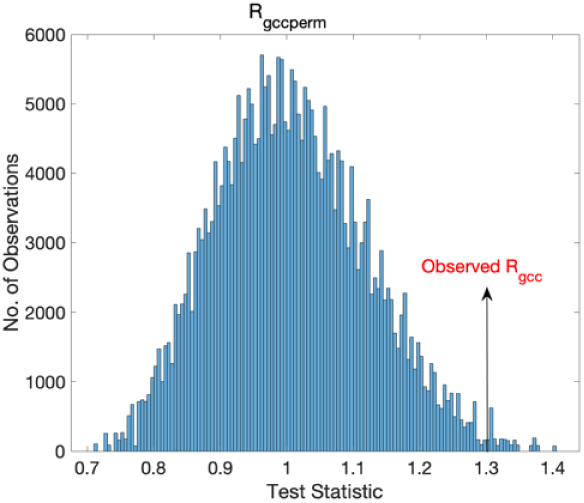
Empirical distribution generated by permutation for GCC analysis on the music state data of both the groups

**Fig. 11.**
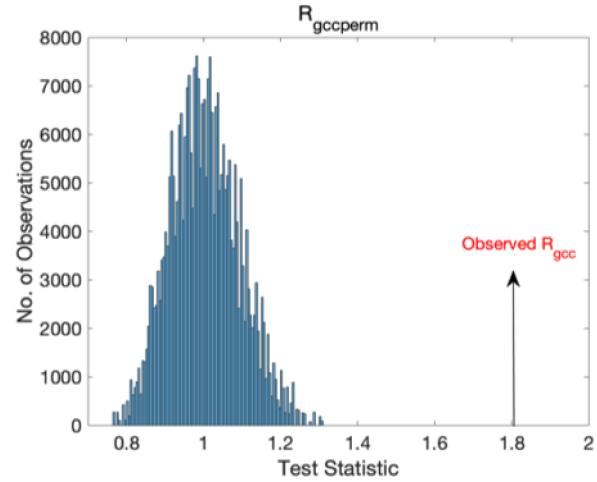
Empirical distribution generated by permutation for GCC analysis on the rest state data of both the groups

This figure demonstrates that the functional brain networks of the two groups under study i.e. the TDC and the ID groups, for rest state data, are indeed statistically different. Also, the lower GCC values for ID population suggests that, perhaps, these subjects have lesser coordination or connectivity strength within their networks. Overall, the functional brain networks of both the groups are statistically different.

## V. CONCLUSION

This study has made some important contributions. First, the data of ID subjects was captured using 14-channel dry electrode EEG system. Although multiple channel wet electrode EEG systems are considered more reliable compared to the dry electrode systems with limited channels, it was worthwhile to test dry electrode system in this study because ID subjects accepted this headset as a playful device and hence, captured data was reliable as proved via statistical analysis as well as via cognitive corroborations of the extracted networks. Since, in general, these subjects do not accept any device or equipment easily, particularly, the gel-based wet electrode system could have disturbed their natural brain response because of anxiety and discomfort. Thus, this study establishes the utility of dry electrode based limited channel EEG devices that can make data collection and analysis of EEG related studies much easier on the ID population.

Second, this work studied functional brain networks formed during the music and rest states of human brain using tensor-based signal processing framework. The task or activity specific functional brain network and functional connectivity of subjects were successfully extracted from their EEG recordings. Distinctions were observed between the rest and music states of the brain by two major analyses, namely, network analysis and network parameter analysis. In the network analysis, the associated BAs and their cognitive functionalities were utilized to reason out the resulting networks. Overall, lesser brain activation was observed in ID group as compared to TDC group, reflecting lesser involvement in executive processing of the task. However, improvement in activation by a cognitive training exercise can presumably help in improving their mental functioning depending on the focus of the cognitive training. This may also be reflective of the benefits observed in population of individuals with traumatic brain injury receiving specific cognitive training for improving a particular cognitive deficit.

Third, in the network parameter analysis, GCC, a network parameter was studied extensively. This parameter analysis was based on the hypothesis that although brain of ID subjects perceives stimuli and show activations, the intra-community strength may be lower in ID population. Indeed, this was observed through the permutation testing framework that was extended to GCC coefficient in this work. Overall, three statistical measures, namely ranksum test, boxplots, and a novel approach of permutation testing framework were utilized to establish the differences in the networks in music versus rest states as well as in ID versus TDC population.

Fourth, the insignificant differences within the groups in music vs. rest states suggests that the preservation of the functional circuits remains more or less intact within the healthy or ID population. However, there is an altered network architecture in different population groups as suggested by the observed group differences. Furthermore, functional networks suggested differential activation of networks during the rest and music states among the groups suggesting poorer attention and lesser involvement in executive planning along with poorer integration of cortical circuitry requiring more neuronal efforts in ID group participants. This may explain the cognitive deficits and associated social impairment observed in such individuals.

In future, researchers should focus on identifying responses to different varieties of environmental stimulation in this population to gather more information regarding the brain networks. The information can then be utilized in developing programs to foster targeted intervention towards environmental manipulation of various stimuli to bolster cognitive rehabilitation in ID population. Specifically, from a practical perspective, training modules focused on strengthening the neural networks of task-associated learning can also help in building the abilities in the ID population. It has been found that task-specific functioning can be strengthened by utilizing neuro-modulation techniques such as transcranial direct current stimulation (tDCS), applied while performing a cognitive task. Motivated with the above, this study suggests to devise interventions utilizing the methods and finding of this work along with the tDCS based design to help ID individuals facing challenges in translating the required actions like performing a task while listening to the instructions simultaneously. A similar strategy can be applicable in this population group to understand how instruction-linked brain activation becomes a helping factor for quick motor-sensory response for the task.

## ACKNOWLEDGMENT

The authors wish to thank Aakash Deep for his support in the data collection and Venkata Ratnadeep Suri for his insightful suggestions on this work.

