## Supplementary for "Studying functional brain networks from dry electrode EEG set during music and resting states in neurodevelopment disorder"

### I. Brodmann Area

Fig. S1 depicts the parcellation of brain into different Brodmann regions. The mapping of Brodmann regions to EMOTIV electrodes along with respective cognitive functionalities is depicted in Table-S1.

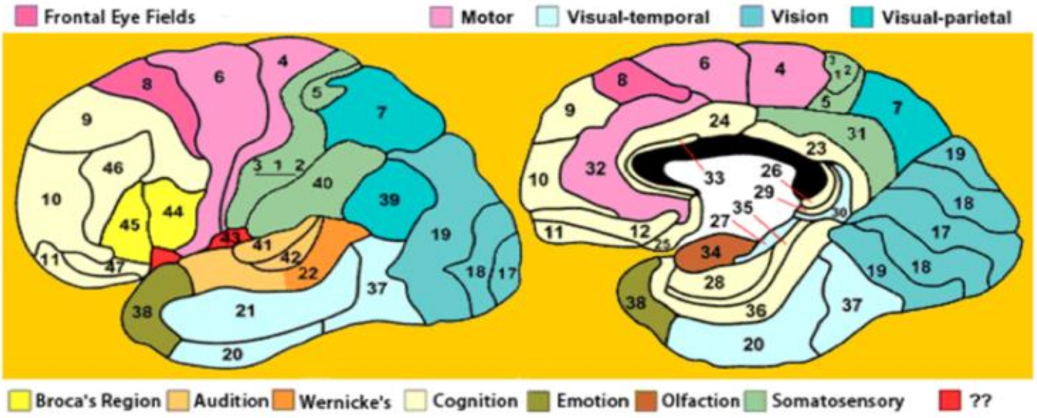


Fig. S1. Brodmann Regions
Source: Adapted from [1]

### II. Table S1 Mapping of Brodmann areas with Emotiv electrodes and their cognitive functionality [2]

| Electrode Name | Brodmann Area | Cognitive functionality |
|---|---|---|
| AF3<br>AF4 | BA09 | Language comprehension and expression, working memory, memory retrieval and encoding, selective attention to sounds, executive planning, auditory imagery and emotional stimuli. |
| F7<br>F8 | BA45<br>BA47 | Language comprehension and expression, working and episodic memory, music enjoyment and attention to speech. |
| F3<br>F4 | BA08 | Secondary motor response, motor planning, imaging and learning, auditory imagery, working memory and visio-motor response. |
| P7<br>P8 | BA37<br>BA19 | Facial detection, experiencing and processing emotions, deductive reasoning, language-oriented tasks and semantic categorization. |
| FC5<br>FC6 | BA06 | Planning of complex, coordinated movements, attention to human voices. Mirror neurons are also present here. |
| T7<br>T8 | BA21 | Basic and complex auditory processing, language comprehension, prosody (rhythmic patterns) comprehension and attention to speech. |
| O1<br>O2 | BA18 | Saccadic movements, visual perception (light, intensity, pattern), visual mental imagery and visual attention. |

### III. Tables for Group level statistical analysis

Table S2 Group level statistical comparisons for GCC-1 and GCC-2 for TDC and ID

| Comparison | GCC-1 | | | GCC-2 | | |
|---|---|---|---|---|---|---|
| | TDC (median) | ID (median) | P-value | TDC (median) | ID (median) | P-value |
| TDC vs ID Rest | 0.4049 | 0.3071 | 3.29E-51 | 0.4215 | 0.3000 | 3.10E-86 |
| TDC vs ID Music | 0.4168 | 0.3020 | 7.84E-106 | 0.4322 | 0.2908 | 1.88E-184 |

Table S3 Group level statistical comparisons for GCC-1 and GCC-2 for Rest and Music

| Comparison | GCC-1 | | | GCC-2 | | |
|---|---|---|---|---|---|---|
| | Rest (median) | Music (median) | P-value | Rest (median) | Music (median) | P-value |
| Rest vs Music TDC | 0.4049 | 0.4168 | 0.0422 | 0.4215 | 0.4322 | 0.0237 |
| Rest vs Music ID | 0.3071 | 0.3020 | 0.1232 | 0.3000 | 0.2908 | 0.0231 |
